# Heterogeneity of cognitive decline in dementia: a failed attempt to take into account variable time-zero severity

**DOI:** 10.1101/060830

**Authors:** Steven J Kiddle, Alice Parodi, Caroline Johnston, Chris Wallace, Richard JB Dobson, for the Alzheimer’s Disease Neuroimaging Initiative (ADNI), for the Australian Imaging Biomarkers and Lifestyle flagship study of ageing (AIBL), and for the Coalition Against Major Diseases (CAMD)

## Abstract

**NOTE:** The biases seen in simulations of Temporal Clustering appear to be even worse in real applications. While this was a novel and interesting approach, ultimately this work has been discontinued. I feel the biases are due to propagated error in estimating individual level offsets based on a single noisy measure, amplified by the fact that high MMSE scores change very slowly and therefore many estimates are from near an asymptote on the left hand of the model. We continue to work in this area, with other approaches showing significantly more promise.

Understanding heterogeneity in Alzheimer’s disease (AD) progression is critically important for the optimal design of trials, allowing participants to be recruited who are correctly diagnosed and who are likely to undergo cognitive decline. Current knowledge about heterogeneity is limited by the paucity of long-term follow-up data and methodological challenges. Of the latter, a key problem is how to choose the most appropriate ‘time zero’ to use in longitudinal models, a choice which affects results. Rather than a pre-specified ‘time zero’ we propose a novel methodology – Temporal Clustering – that defines a new ‘time zero’ using individual offsets inferred from the data. We applied this to longitudinal Mini-Mental State Examination (MMSE), where this approach ensures that individuals have similar estimated MMSE scores at this new ‘time zero’. Simulations showed that it could accurately predict cluster membership after the application of a filter. Next we applied it to a cohort of 2412 individuals, with large variability in MMSE score at first visit. Temporal Clustering was used to split individuals into two clusters. The group showing faster decline had higher average levels of AD risk factors: cerebrospinal fluid tau and *APOE ∊*4. Cluster membership predicted by Temporal Clustering was less affected by individuals’ cognitive ability at first visit than was the case for clusters found using Latent Class Mixture Models. Further application and development of this method will help researchers to identify risk factors affecting cognitive decline.

## 1 Introduction

Patient heterogeneity in cognitive decline can be modelled to aid in the design of clinical trials for AD [1]. The current FDA-approved model uses a generalised logistic (i.e. sigmoidal) curve and includes only baseline cognitive ability, age, gender and the *APOE ∊*4 genetic risk allele [2]. Extensions to the FDA-approved model taking into account additional factors should allow improved design of clinical trials. Additionally, heterogeneity in cognitive decline may be indicative of dementia type, and may therefore help to identify individuals who have been misdiagnosed.

Unfortunately, our understanding of heterogeneity in cognitive decline and the risk factors that affect it are limited. It is challenging to study because of short-term follow-up of individuals in most existing studies. For example, cohort studies such as Alzheimer’s Disease Neuroimaging Initiative (ADNI) [3] and clinical trials such as those in the Coalition Against Major Diseases (CAMD) [4] have relatively short-term follow-up over months or a few years, while the course of cognitive decline in individual patients can take a decade or more [5].

As well as having limited temporal resolution, studies of cognitive decline are further complicated by the fact that individuals can be recruited at different times relative to onset of symptoms. A consequence of this is that it can be hard to choose the most appropriate ‘time zero’ for longitudinal modelling. Commonly used ‘time zero’ options include a fixed age, dementia diagnosis, death or first visit, but this choice can have large effects on the results and replicability of analyses [6, 7].

For a given ‘time zero’, a standard approach to analyse rate of change in cognitive ability is the linear mixed model, which include both fixed effects (i.e. covariates) and random effects. This approach has been applied to identify risk factors of cognitive decline, but has been shown to lead to inflated type 1 error rates and confusion between the effect of risk factors on mean level and rates of change [8].

Extensions to the linear mixed model perform better, but are still limited in their ability to model variability in cognitive ability at ‘time zero’, however defined. For example, Latent Class Mixture Models (LCCMs) extend mixed models to include ‘classes’, i.e. groups of individuals who follow similar trajectories over time. These classes are ‘latent’ (i.e. not known in advance) and are therefore inferred from the data. As such they are a natural framework to summarise heterogeneity in cognitive decline. LCMMs have been used to find clusters of individuals with different trajectories of cognitive decline who differ according to psychotic symptoms [5] or time to a dementia diagnosis [9]. However, in some applications the trajectories appear to differ mostly in individuals’ scores at ‘time zero’ (i.e. disease stage), which is problematic as it may not reflect distinct trajectories (e.g. [5]).

Other researchers have taken a different approach, estimating a random change point that can be seen as an inferred ‘time zero’ (e.g. [10, 11]). These assume that cognitive decline can be divided into two linear sections, representing normal decline and accelerated decline due to dementia respectively. The change point at which they meet is learned from the data. However, the assumption of linearity before and after the change point is probably unrealistic which may bias results.

Rather than use a change point model, other researchers have assumed sigmoidal (i.e. logistic) or exponential trajectories of cognitive decline, with individual offsets (a random effect) used to align each individual to the trajectory model [12, 13]. These models have been used to study the temporal relationship of different biomarkers, leading to models that are only partially consistent with the theoretical model of Jack Jr et al. [14]. Neither of these approaches has been used to develop a clustering approach to help identify risk factors affecting rates of cognitive decline. Outside of dementia research joint clustering and alignment methods have been introduced [15, 16, 17], but these methods depend on densely sampled regular timepoints, which are not available on cognitive decline.

In this work we aimed to study heterogeneity in cognitive decline using data with large ‘time zero’ variability, a problem that is particularly acute when studying disease progression using Electronic Health Records [18, 19]. However, to allow others to verify our work we used data from open access traditional cohort studies in this analysis, combined so that the total dataset has large ‘time zero’ variability. We applied a novel method which combines the benefits of clustering and an inferred ‘time zero’ to improve the analysis of longitudinal heterogeneity. Further, we compare the ability of Temporal Clustering and LCMMs to generate clusters that are related to AD risk factors. To our knowledge this is the second application of such a method to study disease progression [18], and the first attempt to use this type of approach to study cognitive decline. Furthermore, all analyses are done with open source code to allow our work to be verified and extended by others.

## 2 Methods

All analysis in this paper has been performed in R [20], scripts are freely available at https://github.com/KHP-Informatics/TC.

### 2.1 Datasets

We combined data including time series of MMSE from two prospective cohort studies, ADNI1 [3] and the Australian Imaging, Biomarkers and Lifestyle study of aging (AIBL) [21], as well as the placebo arms of the two longest clinical trials from the Coalition Against Major Diseases consortium (CAMD; C-1013 and C-1014 [4]). Studies were combined to make a large dataset, all with short follow-up, but with a lot of variability in baseline cognitive ability. They could be combined as MMSE was recorded consistently across all studies. Individuals were classified as having Normal Cognition (NC), Mild Cognitive Impairment (MCI) or AD. In CAMD, where diagnoses were not given, they were inferred based on MMSE in the following way: MMSE 24 – 30 = NC, MMSE 18 – 23 = MCI, or MMSE 0 – 17 = AD [22]. For more details see Supplementary Methods.

For ADNI and AIBL, genomic DNA was extracted from whole blood with *APOE* genotyped using either TaqMan probes for Single Nucleotide Polymorphisms (rs429358, rs7412) or the *Hha1* restriction enzyme, and assessed using Polymerase Chain Reaction [23, 24]. Levels of total tau in cerebrospinal fluid from ADNI participants have been measured using the xMAP Luminex platform, and were downloaded as ‘UPENNBIOMARK2.csv’ (http://www.adni-info.org/).

### 2.2 Temporal Clustering

Full technical details are provided in Supplementary Methods and are summarised here. For Temporal Clustering, a parametrised trajectory curve (*ϕ*(*t*; *θ*)) has to be supplied, where *t* is time and *θ* is a vector of trajectory parameters. For this application we first used a three parameter sigmoidal trajectory (Supplemental Methods), based on the assumptions of Jack Jr et al. [14], but we found that a two parameter exponential decline curve fitted equally well (*ϕ*(*t*; *θ*):= *θ*_1_ *− exp* (*θ*_2_*t*)), and we therefore used this for inference as it requires fewer parameters. Here, *θ*_1_ represents the maximum MMSE of the trajectory model and *θ*_2_ is an exponential decline rate.

The Temporal Clustering model consists of trajectory parameters (*θ^k^*) for each cluster (*k*), and offsets (*δ_i_*) for each individual (*i*). These offsets are used to shift time points to better align individuals to cluster trajectories, i.e. *ϕ*(*t*+*δ_i_*; *θ^k^*) is the expected MMSE score at timepoint *t* for individual *i* in cluster *k* (Figure 1). Using the exponential decline trajectory, the offset 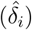 for an individual (*i*) is an estimate of the time between first MMSE assessment and the time at which their MMSE score reached one MMSE point below the maximum MMSE of the model, i.e. an MMSE of *θ*_1_ *−*1.

**Figure 1.**
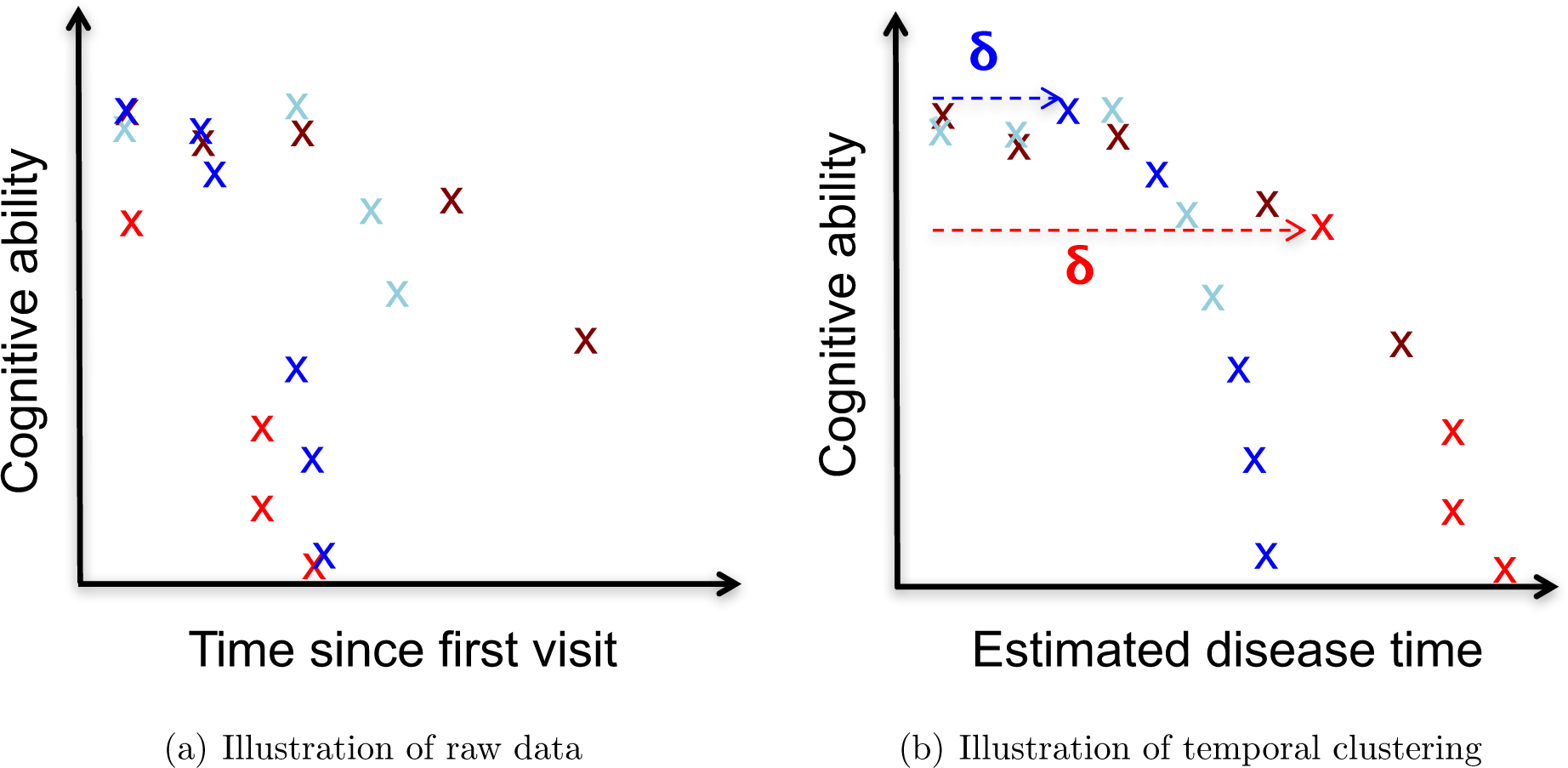
Illustration of (a) raw longitudinal cognitive ability data and (b) the results of applying temporal clustering (*K* = 2). Different individuals are represented by different colours, with different shades of the same colour used to indicate individuals following the same decline trajectory after alignment. Solid curves in (b) are used to represent the long-term trajectory models (*ϕ*(*t*;*θ^k^*)) for the blue and red clusters. The individual offsets (*δ*) applied to dark blue and light red are indicated by dashed lines.

A simplifying assumption of Temporal Clustering is that the baseline parameter (*θ*_1_) takes the same values across all clusters, this was found to be necessary to get good performance and provide identifiability given short follow-up (Supplementary Results).

Temporal Clustering is based on *K*-means, a commonly used clustering algorithm. *K*-means finds clusters by initially allocating all individuals to *K* clusters at random, and then iterating two steps until the model converges. In *K*-means step (1) involves calculating a mean point over all cluster members, and step (2) involves re-allocating individuals to the cluster whose mean they are closest to.

The difference between *K*-means and Temporal Clustering is that rather than calculating cluster means in Step (1), instead trajectory parameters 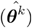 are inferred for each cluster (*k*), along with individual offsets 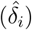. This iterates with cluster re-assignment (Step 2) until convergence.

In this study, for simplicity, we will seek to split individuals into a groups of slower and relatively faster decliners (i.e. *K* = 2). A discrimination score was calculated to assess the relative quality of fit of each individual’s data to their assigned versus unassigned cluster (Supplemental Methods).

### 2.3 Simulation study

Full details of the simulation study are available in Supplementary Methods but are briefly summarised here. We performed simulations based on the combined cohort structure, i.e. with the same number of individuals, similar time points and with longitudinal MMSE data resembling the real data when plotted. A range of true parameters *θ* were used, with 100 simulations performed for each set. For each simulation true offsets *δ_i_* were generated at random, and timepoints assigned to simulate missing data due to death (i.e. missing data when MMSE ≤ 0). The accuracy of clustering was assessed using the Adjusted Rand Index (ARI), which takes values from zero (i.e. no better than chance) to one (i.e. perfect clustering). ARI was calculated using the R package ‘mclust’.

### 2.4 Latent Class Mixture Models

LCMMs were performed using the ‘lcmm’ package using spline models with default parameters and no covariates (i.e. no fixed effects). ‘Time zero’ was chosen to be either first visit or 50*^th^* birthday. As in Lima-Proust et al. [9] LCMMs were run on normMMSE, that is MMSE scores transformed to have better metric properties [25]. For comparison we also ran LCMM on raw MMSE scores.

### 2.5 Statistical analysis

Time was coded as days since first visit. We analysed the simulated and real datasets using Temporal Clustering.

Logistic regression was used to build multivariate models of cluster membership using the R package ‘glm’. For regression models baseline MMSE points were recoded in units of ten (i.e. divided by 10), and cerebrospinal fluid tau was recoded in units of 100 pg/ml (i.e. divided by 100). Gender, cohort and the number of *APOE ∊*4 alleles were coded as categorial variables. CAMD individuals with an age *>* 89 years have had their age recorded as 999 to reduce identifiability, these 13 individuals were excluded from regression analyses. Individuals with missing *APOE* or cerebrospinal fluid tau data were excluded from the relevant analyses.

‘MMSE at first visit’ analysis included covariates: age at first visit, gender and cohort. *APOE* analyses were performed on ADNI1 and AIBL individuals and included co-variates gender and cohort as well as age and MMSE at first visit. Cerebrospinal fluid tau analyses included co-variates gender, and the number of *APOE ∊*4 alleles as well as age and MMSE at first visit.

## 3 Results

Demographic information on the individual and combined cohorts are provided in Table 1. The distribution of MMSE differs between these cohorts, generally individuals in the CAMD have lower MMSE scores at first visit (Figure S1). As was our intention, combining these cohorts therefore results in a large variability in MMSE at first visit (interquartile range (15 – 27), Figure 2(a)).

**Table 1.** Demographic summary of cohorts. Diagnosis for ADNI and AIBL has been recorded, whereas for C-1013 and C-1014 it has been inferred from MMSE scores. Data is summarised using counts or median (interquartile Range). TS = time series, ADNI = Alzheimer’s Disease Neuroimaging Initiative, AIBL = Australian Imaging Biomarkers and Lifestyle flagship study of ageing, MMSE = Mini-Mental State Examination, NC = Normal Cognition, MCI = Mild Cognitive Impairment, AD = Alzheimer’s disease, and *APOE ∊4* = the *∊*4 allele of the *apolipoprotein E* gene.

| Cohort | Time series with > 1 time point | Number of time points | Length of time series in years | Diagnoses at first visit (NC/ MCI/ AD) | Gender (Female /Male) | Age at first visit in years | Number of $APOE \epsilon_4$ alleles (0/1/2) |
| --- | --- | --- | --- | --- | --- | --- | --- |
| Combined | 2412 | 5 (4 - 6) | 1.9 (1.5 - 3.1) | 399/920/1093 | 1186/1226 | 75 (70 - 80) |  |
| ADNI | 785 | 5 (4 - 6) | 3.1 (2.1 - 3.2) | 222/382/181 | 327/458 | 75 (70 -79) | 397/301/87 |
| AIBL | 264 | 3 (3 - 4) | 4.4 (3.1 - 4.6) | 177/52/35 | 136/128 | 72 (66 - 79) | 135/110/19 |
| C-1013 | 719 | 5 (5 - 6) | 1.6 (1.4 - 2.1) | . /247/472 | 361/358 | 76 (70 - 80) | Unknown |
| C-1014 | 644 | 5 (4 - 5) | 1.5 (0.68 - 1.6) | . /239/405 | 362/282 | 76 (70 - 81) | Unknown |

**Figure 2.**
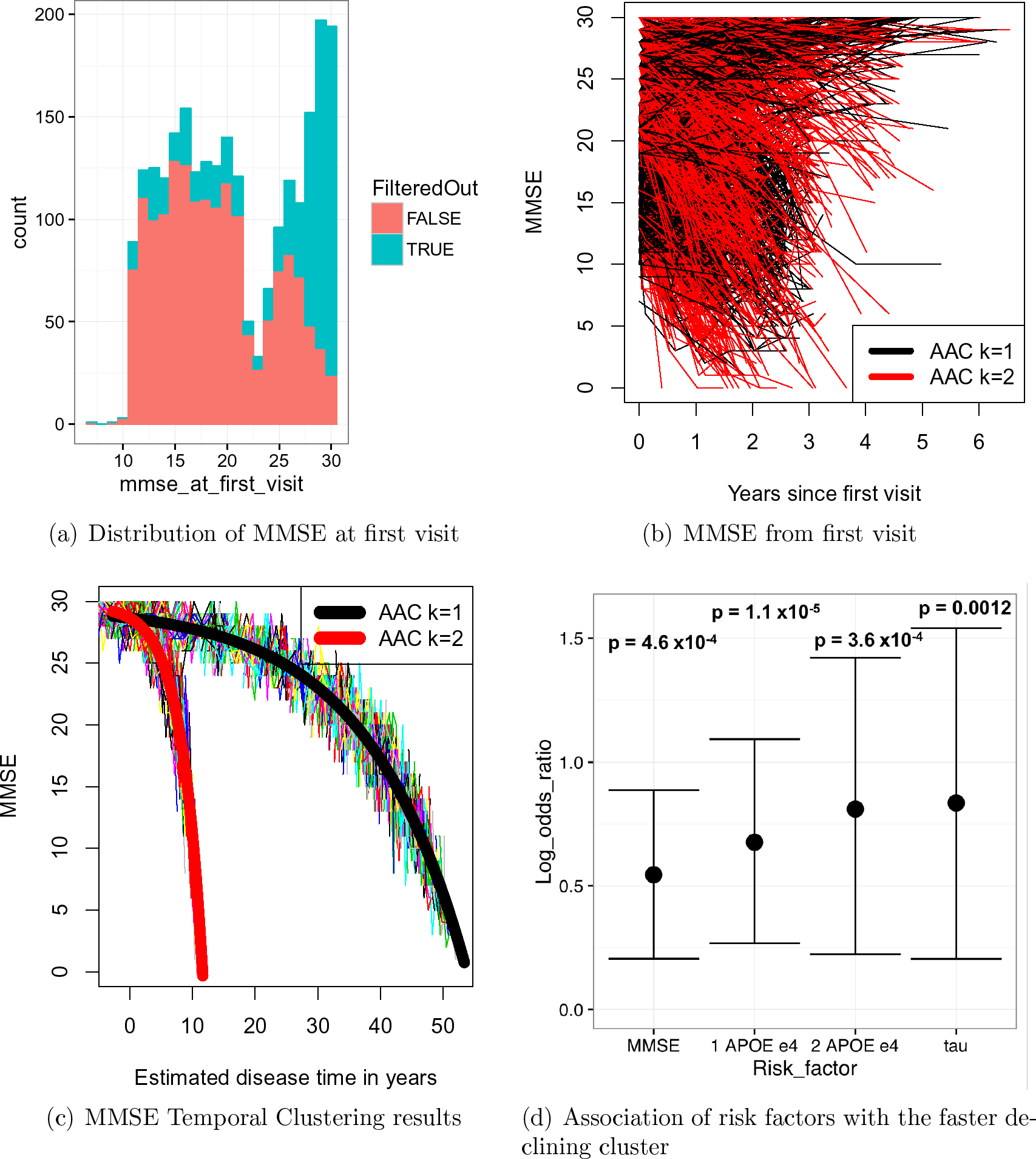
(a) Stacked histogram showing the distribution of MMSE at first visit for the combined cohort (both colours), with the distribution after applying a discrimination score threshold of 2 shown in pink. Spaghetti plots of longitudinal MMSE in the combined cohort with (b) first visit as ‘time zero’ or (c) a ‘time zero’ inferred by Temporal Clustering. Unfiltered clusters (*k*) are represented by either (b) the colour of individual curves or (c) the colour of cluster trajectories (represented by thick curves). (d) Log odds ratios and confidence intervals for risk factors (MMSE at first visit in units of ten, number of *APOE ∊*4 alleles or cerebrospinal fluid tau levels in units of 100 pg/ml). 29

Longitudinal MMSE for individuals with first visit as ‘time zero’ is shown in Figure 2(b), where a large amount of heterogeneity can be seen. The 2,412 individuals in the combined cohort were followed-up over a median of 1.9 years at a median of 5 visits. At first visit 42% had AD (diagnosed or inferred), whereas at the last visit 60% did (Table 1 and Table S1).

### 3.1 Accuracy of clustering: a simulation study

After using simulated data to choose default values for Temporal Clustering parameters (Supplementary Results), we then used it to assess clustering accuracy, i.e. to assess whether Temporal Clustering could accurately distinguish between a slower and faster group of cognitive decliners. With no filter applied, the clustering result was only slightly better than would be expected by chance (ARI 0.12; Figure 3).

**Figure 3.**
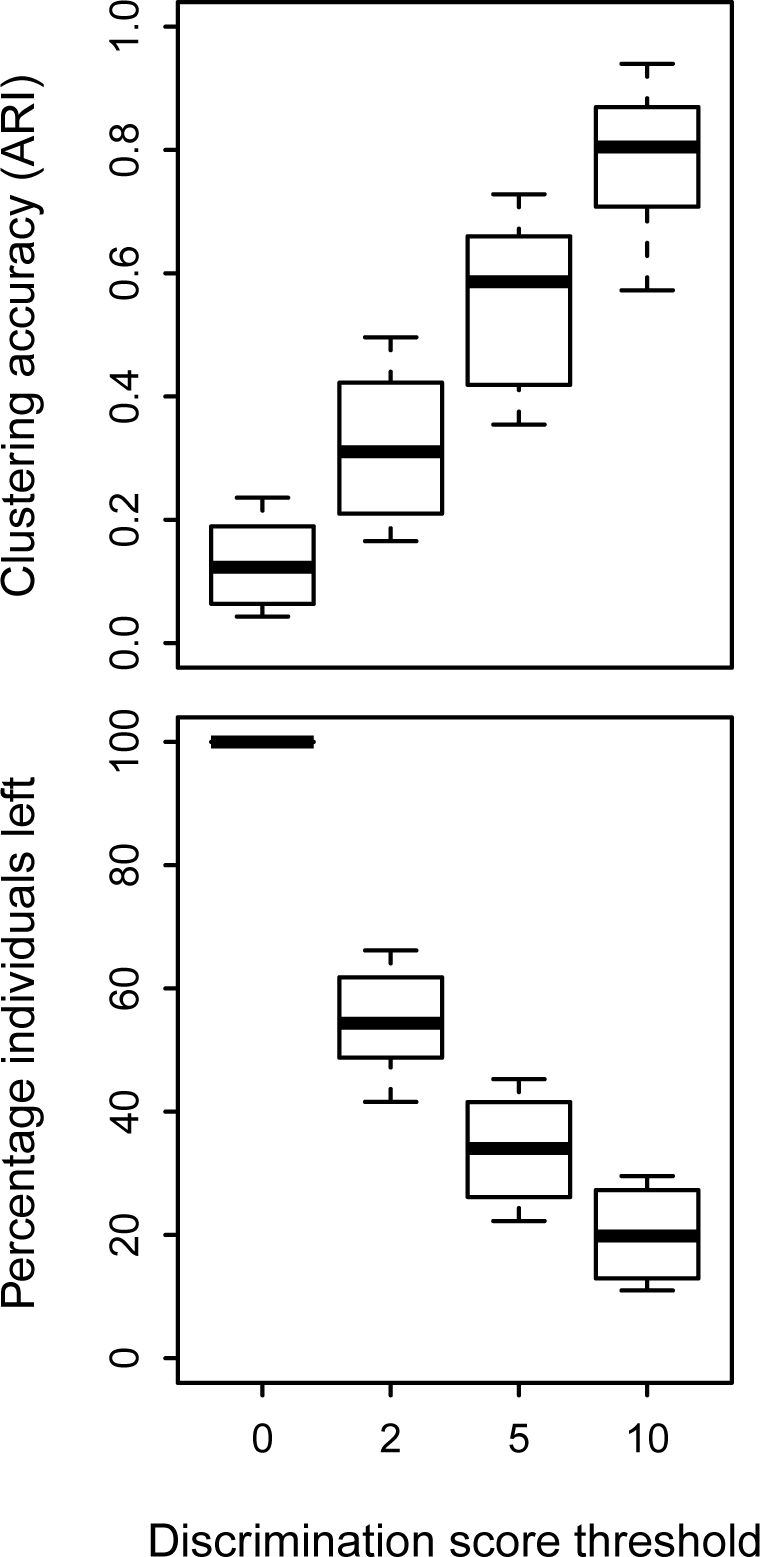
Boxplots of simulation study examining the effect of a discrimination score filter on the accuracy of Temporal Clustering cluster assignment, and number of individuals left after the filter is applied. Discrimination score is measure of relative goodness of fit of each individual to their assigned cluster. Clustering accuracy is measured in Adjusted Rand Index (ARI). Boxplots are over 100 simulations of four different choices of 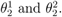

Clustering accuracy increased as individuals were filtered out on the basis of the discrimination score, a measure of confidence in assignment of the individual to a cluster (Figure 3). A tradeoff can clearly be seen where the more stringent the filter (i.e. the higher the threshold) the higher the clustering accuracy and the lower the number of individuals left after filtering. In our application the risk factors of interest, *APOE* and cerebrospinal fluid tau, are only recorded for 43% and 17% of individuals respectively and so we prioritised retaining individuals above clustering accuracy. For this reason we selected a discrimination score threshold of 2, which led to a median clustering accuracy (ARI) of 0.31 but retained over half the individuals (median 54%). A much more stringent threshold of 10 led to a clustering accuracy (ARI) of 0.80, but retained fewer than a fifth of individuals on average.

The most obvious characteristics of simulated individuals removed by a discrimination score filter of 2 were that they had higher MMSE scores at first visit (Figure S2a) and/or less than a year of follow-up (Figure S2b).

### 3.2 Temporal Clustering can distinguish between faster and slower decliners

We next sought to use Temporal Clustering (*K* = 2) to summarise cognitive decline in the combined cohort. Before the discrimination score filter was applied this resulted in one slowly declining cluster containing 1,335 individuals and another which declined faster and contained 1,077 individuals (Figure 2(c)). The estimated maximum MMSE 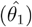 of the model was 30. The estimated yearly rate of exponential decline in MMSE score was 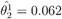 for the slowly declining cluster and 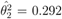 for the faster declining cluster. The trajectory of the faster declining cluster takes approximately 10 years to go from an MMSE of 29 to 0, which fits well with LCMM trajectories learned from AD patients [5], whereas the more slowly declining trajectory is estimated to take approximately 50 years (Figure 2(c)).

As our simulation study suggested it was a sensible step, we filtered out individuals for who we didn’t have sufficient confidence in cluster assignment (i.e. those with a discrimination score *<* 2). This resulted in the removal of 31% of individuals, leaving 969 members (73%) of the slowly declining cluster and 688 members (64%) of the faster declining cluster. As in the simulation study, the filter removed a greater number of individuals with higher MMSE at first visit, presumably as the differences between clusters at that stage are more subtle (Figure 2(a)).

We examined diagnosis at last visit for individuals in ADNI1 or AIBL who remained after the filter, as these were the only cohorts with diagnostic information provided (instead of inferred). Approximately half of these individuals in the slow declining cluster had AD at last visit (99/191), in comparison to *∼*80% (236/301) in the faster declining cluster.

### 3.3 Association between AD risk factors and AD-like cognitive decline

We next sought to use the filtered Temporal Clustering results to identify risk factors distinguishing the two clusters. Higher MMSE at first visit was associated with membership of the faster declining cluster (log odds ratio (*LOD*) for 10 point increase = 0.54, p-value = 1.7×10^*−*3^). Focusing on the subset of individuals with *APOE* data, we showed a positive and dose dependent association between the number of *APOE ∊*4 alleles and membership of the faster declining cluster (1 allele *LOD* = 0.67, p-value = 1.3 *×* 10^*−*3^; 2 alleles *LOD* = 0.81, p-value = 7.8 *×* 10^*−*3^). Finally, we found an association between the fast declining cluster and the level of cerebrospinal fluid tau at first visit in ADNI (*LOD* for 100 pg/ml increase = 0.84, p-value = 0.014). The association of first visit MMSE, *APOE* and tau with the faster declining cluster is visualised in Figure 2(d). Results were consistent when no filter was used (Table 2).

**Table 2.** Table summarising logistic regression analysis, comparing cluster membership to AD risk factors for Temporal Clustering (TC) and LCMM. Four different LCMM models have been run, combining one of two choice for ‘time zero’ with the choice to use raw MMSE or normalised MMSE (normMMSE). Each line refers to a different logistic regression analysis to better cater for missing risk factor data, except for *APOE* for each clustering method, which were modelled together. Signs for *LOD* have been swapped when appropriate to allow appropriate comparisons, as signs depend on cluster labels which can be swapped arbitrarily. MMSE at first visit is coded in units of ten and cerebrospinal fluid tau is coded in units of 100 pg/ml. N = sample size with complete data, *LOD* = Log Odds Ratio, SE = Standard Error, LCMM = Latent Class Mixture Model, MMSE = Mini Mental State Examination, CSF = cerebrospinal fluid.

| Method | Time zero | Variable | Risk factor | N | *LOD* | SE | p-value |
| --- | --- | --- | --- | --- | --- | --- | --- |
| TC | Inferred | MMSE | MMSE first visit | 2399 | 0.5 | 0.14 | $4.6 \times 10^{-4}$ |
| TC | Inferred | MMSE | 1 APOE e4 | 1049 | 0.61 | 0.14 | $1.1 \times 10^{-5}$ |
| TC | Inferred | MMSE | 2 APOE e4 | 1049 | 0.84 | 0.23 | $3.6 \times 10^{-4}$ |
| TC | Inferred | MMSE | CSF tau | 407 | 0.77 | 0.24 | 0.0012 |
| LCMM | First visit | MMSE | MMSE first visit | 2399 | -1.6 | 0.23 | $1.3 \times 10^{-11}$ |
| LCMM | First visit | MMSE | 1 APOE e4 | 1049 | 0.4 | 0.33 | 0.22 |
| LCMM | First visit | MMSE | 2 APOE e4 | 1049 | 0.64 | 0.41 | 0.12 |
| LCMM | First visit | MMSE | CSF tau | 407 | 0.38 | 0.4 | 0.35 |
| LCMM | Age 50yrs | MMSE | MMSE first visit | 2399 | -1.7 | 0.25 | $3.5 \times 10^{-12}$ |
| LCMM | Age 50yrs | MMSE | 1 APOE e4 | 1049 | 0.17 | 0.34 | 0.62 |
| LCMM | Age 50yrs | MMSE | 2 APOE e4 | 1049 | 0.31 | 0.45 | 0.49 |
| LCMM | Age 50yrs | MMSE | CSF tau | 407 | 0.49 | 0.4 | 0.21 |
| LCMM | First visit | normMMSE | MMSE first visit | 2399 | -10 | 0.72 | $5.0 \times 10^{-48}$ |
| LCMM | First visit | normMMSE | 1 APOE e4 | 1049 | 0.68 | 0.27 | 0.013 |
| LCMM | First visit | normMMSE | 2 APOE e4 | 1049 | 1.1 | 0.36 | 0.0017 |
| LCMM | First visit | normMMSE | CSF tau | 407 | 1.2 | 0.37 | 0.0015 |
| LCMM | Age 50yrs | normMMSE | MMSE first visit | 2399 | -10 | 0.7 | $3.7 \times 10^{-47}$ |
| LCMM | Age 50yrs | normMMSE | 1 APOE e4 | 1049 | 0.62 | 0.27 | 0.022 |
| LCMM | Age 50yrs | normMMSE | 2 APOE e4 | 1049 | 1.1 | 0.35 | 0.0028 |
| LCMM | Age 50yrs | normMMSE | CSF tau | 407 | 1.1 | 0.36 | 0.0023 |

### 3.4 Temporal Clustering produces clusters that are more significantly associated with *APOE* status than LCMM

We wished to compare the results of Temporal Clustering and LCMMs, specifically which approach produced clusters with higher associations to known AD risk factors (*APOE* and tau). To make the comparison more straightforward, this was performed on Temporal Clustering results before filtering, and on LCMM models without any co-variates. To make it a fairer comparison for LCMM we used two different choice of ‘time zero’ (first visit or 50th birthday) and both raw and normalised MMSE (normMMSE).

Because Temporal Clustering infers a ‘time zero’ we would expect by design its clusters to be less affected by MMSE at first visit than approaches like LCMM that do not. This was indeed the case, confounding of unfiltered Temporal Clustering clusters by MMSE at first visit was at least three-fold lower in absolute terms than that achieved by LCMM (*LOD* for a 10 point change = 0.5 for Temporal Clustering versus −1.6 – −10 for LCMM, Table 2). Indeed, MMSE at first visit was by far the most significant predictor of LCMM cluster membership in all cases. The biggest difference in the association of risk factors with unfiltered clusters was for *APOE*, especially the significance of the association of a single *APOE ∊*4 allele with cluster membership (p-value = 1.1 *×* 10^*−*5^ for Temporal Clustering versus 0.62 – 0.013 for LCMM, Table 2).

Overall LCMM cluster membersip only had a clear relationship with AD risk factors when MMSE was normalised, which resulted in a large difference in MMSE at first visit between the clusters (Table 2). In contrast Temporal Clustering cluster membership had a clear relationship to AD risk factors, and a lower difference in MMSE at first visit, with and without a discrimination score filter.

## 4 Discussion

NOTE: The biases seen in simulations of Temporal Clustering appear to be even worse in real applications. While this was a novel and interesting approach, ultimately this work has been discontinued. I feel the biases are due to propagated error in estimating individual level offsets based on a single noisy measure, amplified by the fact that high MMSE scores change very slowly and therefore many estimates are from near an asymptote on the left hand of the model. We continue to work in this area, with other approaches showing significantly more promise.

We have introduced a new method - Temporal Clustering - that can model cognitive decline by combining an estimated individual offset with clustering on that new time-scale. We show that this leads to clusters that are less influenced by MMSE at first visit, which we believe will make it easier to identify risk factors of cognitive decline. To this end we show a dose-dependent enrichment of *APOE ∊*4 carriers in the faster declining (i.e. AD-like) cluster, a difference that is more significant than for clusters produced by LCMM approaches on this dataset.

There is some inconsistency in the literature about the relationship between *APOE ∊*4 and the rate of cognitive decline. Some studies in non-demented individuals have found no relationship, e.g. Winnock et al., [26]. However, the majority of studies have either found a modest relationship (e.g. [27, 28, 29, 30]), or one that depends on other factors such as amyloid beta [31], alcohol [32] and body mass [33]. The inconsistency of these studies may be explained by cohort differences and/or the strong methodological challenges of the study of cognitive decline [7].

In the field of cluster analysis determination of the ‘optimal’ number of clusters is known to be tricky. For example, Bauer et al. [34] have argued that the optimal number of clusters in a model do not necessarily respond to the number of ‘real’ subgroups in an application. Instead they argue that clusters can equally well be interpreted as having no meaning beyond being a convenient summary of non-gaussian distributions. Therefore in this study, for simplicity, we have generated models with just two clusters (*K* = 2). By comparing cerebrospinal fluid tau and *APOE* genotype between clusters we showed that it is plausible that clusters summarise genuine heterogeneity.

As with the choice of any ‘time zero’ other than time in study, Temporal Clustering relies on the convergence assumption, that between-person differences over time are representative of within-person changes [6, 35]. Other assumptions underlying Temporal Clustering include symmetric and independent distribution of errors, as implied by the use of least squares estimation. The bounded nature of MMSE, which takes a minimum of zero and a maximum of 30, means that the true distribution cannot be symmetric. In addition to this Temporal Clustering assumes that data is missing completely at random, a stronger and less realistic assumption than the missing at random assumption of mixed models. However, even with these limitations it is encouraging that reasonable clustering accuracy was achieved in the simulation study after filtering for discrimination score, especially as we explicitly simulated missing data due to death.

While simulations showed the effectiveness of Temporal Clustering at estimating cluster membership, they also showed biased estimation of trajectory parameters, especially for clusters with a slow rate of decline (Supplementary Results). Reducing this bias could be useful in its own right, but may also improve the assignment of individuals to clusters. This bias could be due to overfitting of the individual offsets (*δ_i_*), which could be reduced in the future by penalising unrealistic offsets. Alternatively it could be due to the mixture of between and within individual progression in the model, and the large extrapolation beyond the length of follow-up available (from *∼* 2 years to 10 or even 50 years). Therefore, it is hard to know whether the *∼* 50 year trajectory of the slowly declining cluster truely reflects within-individual change, or is an artefact of the model. This could be tested in a datasets with longer individual follow-up.

Despite the biased estimation of trajectory parameters, the more slowly declining cluster is still striking. From a clinical point of view this trajectory appears to decline too slowly to represent AD. Backing this up is the fact that it does include a higher proportion of individuals with NC or MCI at last visit (Table S1). However, the fact that around half of this cluster have a diagnosis of AD at last visit could suggest a problem with the model, perhaps motivating additional clusters. A less likely alternative hypothesis would be that individuals with a diagnosis of AD in the slowly declining cluster are misdiagnosed.

An advantage of mixed or random change point models over Temporal Clustering is that co-variates can be explicitly modelled, rather than considered post-hoc. Extending Temporal Clustering to consider co-variates could allow it to have more flexibility in the baseline of the model, this could get around the current crude assumption that the maximum MMSE in a lifetime is the same for all individuals.

A limitation of this study is the use of MMSE to measure cognitive decline. MMSE is acknowledged to have ceiling and floor effects and to be relatively insensitive to cognitive change before MCI [36]. We concentrated on MMSE within this study as it is one of the most widely collected measures of cognitive ability in dementia. For example, longitudinal MMSE data is available for thousands of patients at the South London and Maudsley NHS Foundation Trust, where it has been extracted from Electronic Health Records from routine care [19]. However, the method should generalise to other measures of cognitive ability.

In conclusion we have demonstrated that it is possible to model cognitive decline using a combination of clustering and inference of individual offsets. This reduces, but does not eliminate, the effect of baseline MMSE on cluster assignment. Finally, we demonstrated a relationship between clusters and known AD risk factors. We believe that Temporal Clustering and future extensions will be useful for studying progression of dementia biomarkers. To allow others to repeat this analysis and explore extensions the source code is freely available at https://github.com/KHP-Informatics/TC.

NOTE: The biases seen in simulations of Temporal Clustering appear to be even worse in real applications. While this was a novel and interesting approach, ultimately this work has been discontinued. I feel the biases are due to propagated error in estimating individual level offsets based on a single noisy measure, amplified by the fact that high MMSE scores change very slowly and therefore many estimates are from near an asymptote on the left hand of the model. We continue to work in this area, with other approaches showing significantly more promise.

## 5 Acknowledgements

SJK is supported by an MRC Career Development Award in Biostatistics (MR/L011859/1). CW is funded by the Wellcome Trust (107881) and the MRC (MC_UP_1302/5).

The author CJ receives salary support and the authors acknowledge infrastructure support from the National Institute for Health Research (NIHR) Biomedical Research Centre for Mental Health. The views expressed are those of the author(s) and not necessarily those of the NHS, the NIHR or the Department of Health.

We would like to acknowledge John Todd and Ken Smith (University of Cambridge), as well as Sylvia Richardson (MRC Biostatistics Unit), for hosting SJK during this work and manuscript review. The JDRF/Wellcome Trust Diabetes and Inflammation Laboratory is in receipt of a Wellcome Trust Strategic Award (107212) and receives funding from the JDRF (5-SRA-2015-130-A-N) and the NIHR Cambridge Biomedical Research Centre. The Cambridge Institute for Medical Research (CIMR) is in receipt of a Wellcome Trust Strategic Award (100140).

The research leading to these results has received support from the Innovative Medicines Initiative Joint Undertaking under grant agreement no 115372, resources of which are composed of financial contribution from the European Union’s Seventh Framework Programme (FP7-/2007-2013) and in-kind contribution from EFPIA companies.

This work was supported by awards to establish the Farr Institute of Health Informatics Research, London, from the Medical Research Council, Arthritis Research UK, British Heart Foundation, Cancer Research UK, Chief Scientist Office, Economic and Social Research Council, Engineering and Physical Sciences Research Council, National Institute for Health Research, National Institute for Social Care and Health Research, and Wellcome Trust (grant MR/K006584/1).

Data collection and sharing for this project was funded by the Alzheimer’s Disease Neuroimaging Initiative (ADNI) (National Institutes of Health Grant U01 AG024904) and DOD ADNI (Department of Defense award number W81XWH-12-2-0012). ADNI is funded by the National Institute on Aging, the National Institute of Biomedical Imaging and Bio-engineering, and through generous contributions from the following: AbbVie, Alzheimer’s Association; Alzheimer’s Drug Discovery Foundation; Araclon Biotech; BioClinica, Inc.; Biogen; Bristol-Myers Squibb Company; CereSpir, Inc.; Eisai Inc.; Elan Pharmaceuticals, Inc.; Eli Lilly and Company; EuroImmun; F. Hoffmann-La Roche Ltd and its affiliated company Genentech, Inc.; Fujirebio; GE Healthcare; IXICO Ltd.; Janssen Alzheimer Immunotherapy Research & Development, LLC.; Johnson & Johnson Pharmaceutical Research & Development LLC.; Lumosity; Lundbeck; Merck & Co., Inc.; Meso Scale Diagnostics, LLC.; NeuroRx Research; Neurotrack Technologies; Novartis Pharmaceuticals Corporation; Pfizer Inc.; Piramal Imaging; Servier; Takeda Pharmaceutical Company; and Transition Therapeutics. The Canadian Institutes of Health Research is providing funds to support ADNI clinical sites in Canada. Private sector contributions are facilitated by the Foundation for the National Institutes of Health (www.fnih.org). The grantee organization is the Northern California Institute for Research and Education, and the study is coordinated by the Alzheimer’s Disease Cooperative Study at the University of California, San Diego. ADNI data are disseminated by the Laboratory for Neuro Imaging at the University of Southern California.

Major AIBL funders include CSIRO Flagship Program, Science Industry Endowment Fund, NHMRC, Dementia CRC, CRC-Mental Health, Alzheimer’s Association, an anonymous Foundation, GE Healthcare, Janssen, Pfizer, Avid Radiopharmaceuticals, Navidea, Merck, and Astra Zeneca.

We would like to thank Graciela Muniz-Terrera, Rob Howard, Steven Hill, Robert Goudie and peer reviewers for constructive advice.

## 6 Conflict of interest

SJK has served on a Roche Diagnostics advisory board on blood biomarkers for AD. RJBD receives grant support from Eli Lilly for a research project. AP and CW declare no conflict of interest.

